# *De novo* transcriptome assembly of the African bullfrog *Pyxicephalus adspersus* for molecular analysis of aestivation

**DOI:** 10.1101/588889

**Authors:** Naoki Yoshida, Chikara Kaito

## Abstract

The molecular mechanisms of aestivation, a state of dormancy that occurs under dry conditions at ordinary temperature, are not yet clarified. Here, we report the first *de novo* transcriptome assembly of the African bullfrog *Pyxicephalus adspersus*, which aestivates for 6-10 months during the hot and dry season. Polyadenylated RNA from tissues was sequenced to 75,320,390 paired-end reads, and the *de novo* assembly generated 101,682 transcripts. Of these, 100,093 transcripts had open reading frames encoding more than 25 amino acids. BLASTx analysis against the Uniprot *Xenopus tropicalis* protein database revealed 64,963 transcripts having little homology with an E value higher than 1E-5 and 8,147 transcripts having no homology, indicating that the African bullfrog has many novel genes that are absent in *X. tropicalis*. The other 28,570 transcripts had homology with an E value lower than 1E-5 for which molecular functions were estimated by gene ontology (GO) analysis and found to contain the aestivation-related genes conserved among other aestivating organisms, including the African lungfish. This study is the first to identify a comprehensive set of genes expressed in the African bullfrog, thus providing basic information for molecular level analysis of aestivation.

## Introduction

Dormancy is a phenomenon by which organisms temporarily decrease their physiologic activity under conditions such as low temperature, drying, and depletion of energy sources, and is observed in many organisms from microorganisms to mammals. In vertebrates, hibernation and aestivation are two common types of dormancies performed under either a low temperature condition or a drying and ordinary temperature condition, respectively. Molecular biologic and physiologic analyses of vertebrate hibernation have been performed for the Siberian chipmunk and Northern leopard frog. In Siberian chipmunks, hibernation is initiated by two cues: a seasonal change in the photoperiod depending on the melatonin level, and a hibernation-specific protein complex induced by cold temperature [1,2]. Energy-related genes are regulated to decrease body temperature and to suppress the contractile activity of cardiac myocytes. In the Northern leopard frog, glucose and glycerol concentrations in the body are increased to confer freeze tolerance, and the mammalian target of rapamycin (mTOR) is activated to prevent muscle atrophy during hibernation [3,4,5]. These molecular and physiologic aspects of hibernation have attracted attention as the potential basis for novel therapeutics against malignant tumors or tissue necrosis.

Aestivation in vertebrates is observed in many amphibian and fish species. Aestivation is similar to hibernation in terms of the reduced metabolic activity, but it is clearly distinguished from hibernation in terms of the environmental conditions; aestivation is performed under dry conditions at ordinary temperatures, whereas hibernation is performed under aqueous conditions at low temperatures [6]. Therefore, the mechanisms of aestivation are assumed to differ from those of hibernation, such as suppression of energy metabolism at an ordinary temperature or tolerance against dehydration. Molecular level analyses of aestivation are important to uncover mechanisms of dormancy that have not been clarified in studies of hibernation, but such studies are scarce at present.

So far, three species that undergo aestivation – the Green striped burrowing frog (*Cyclorana alboguttata*), Couch’s spadefoot toad (*Scaphiopus couchii*), and African lungfish (*Protopterus annectens*) – have been analyzed at the transcriptional level to identify genes whose expression is induced or suppressed during aestivation [7,8,9]. Very little is known about the molecular mechanisms of aestivation, however, including how energy metabolism is suppressed or how tissue damage is minimized in the aestivating organisms.

In the present study, we aimed to identify genes expressed in the African bullfrog, *P. adspersus* using *de novo* transcriptome assembly. The African bullfrog, a species belonging to the family of *Pyxicephalidae*, is a large frog – body size of males is larger than 20 cm and that of females is ∼14 cm. This frog inhabits a savanna area of east Africa, south Africa, and the southern part of central Africa, where the air temperature ranges from 20-30°C throughout the year, and there are pronounced dry and rainy seasons. During the dry season, the frogs burrow underground and form a tough cocoon to reduce evaporative water loss and decrease the respiration rate to less than 10% that during the active stage [10,11]. The aestivation of the frog continues for 6-10 months. Because aestivation of the frog can be artificially induced and breeding methods are established for breeding the frogs as pets, the African bullfrog is a suitable research model animal to investigate the molecular mechanisms of aestivation. Furthermore, the African bullfrog is phylogenetically and geographically distant from the other two aestivating frogs: the Green striped burrowing frog and Couch’s spadefoot toad, which inhabit Australia and North America, respectively. Therefore, studies of the aestivation of the African bullfrog will help to elucidate the molecular mechanisms of aestivation that are conserved among different aestivating frog species. In this study, we identified and characterized the genes of the African bullfrog to provide basic information for investigating the aestivation mechanism.

## Materials and Methods

### Ethics statement

The Animal Care and Use Committee of Okayama University approved this work (Approval number, OKU-2019300). This study was performed in strict accordance with the recommendation of the Fundamental Guidelines for Proper Conduct of Animal Experiment and Related Activities in Academic Research Institutions under the jurisdiction of the Ministry of Education, Culture, Sports, Science, and Technology in Japan, 2006.

### Experimental animal

A captive bred African bullfrog was purchased from a specialty reptile and amphibian store (Hachurui Club, Nakano, Japan). The purchased frog was maintained in a plastic container with coarse sand (5-7 mm diameter) and water. The frog was fed every day with house crickets (*Acheta domestica*) or wax worms (the larvae of the *Galleria mellonella*) for the first 3 weeks, and then fed every day with artificial diets (Samuraijapan, Ibaraki, Japan) for 2 months.

### Isolation of genomic DNA and total RNA

The young adult frog (15 g body mass) was kept without feeding for 36 h. The frog was anesthetized by placing it into crushed ice for 10 min and then dissected on ice. The intestines were separated, the intestinal contents removed in phosphate buffered saline (pH7.4), and the intestines frozen in liquid nitrogen. The other tissues, including inner organs, muscle, and skin, were quickly excised to 3 - 5 mm^3^ (0.1 - 0.3 g) and frozen in liquid nitrogen. The frozen samples were maintained at −80°C.

Genomic DNA was isolated by a standard phenol/chloroform extraction method [12]. Frozen muscle tissues (5 mm^3^, 0.2 g) were submerged in a mixture of 5 ml of phenol/chloroform/isoamyl alcohol (25:24:1) and 5 ml of TE Buffer (10 mM Tris-HCl [pH 8.0], 1mM EDTA). Then, the tissues were homogenized at 10,000 rpm three times for 30 s each with a Polytron benchtop homogenizer (Kinematica, Switzerland). The sample was centrifuged at room temperature at 2300 *g* for 10 min, and the upper aqueous phase (1.0 ml) was transferred to a new tube. The sample was centrifuged at room temperature at 20,400 *g* for 5 min to completely remove the organic solvents. The upper aqueous phase (0.7 ml) was transferred to a new tube, vortexed with an equivalent volume of chloroform, and centrifuged at room temperature for 5 min at 20,400 *g*. The upper aqueous phase (0.5 ml) was transferred to a new tube, mixed with a 2.5-fold amount of ethanol and a 0.1-fold amount of 3M sodium acetate, and centrifuged at 20,400 *g* for 15 min at room temperature. The precipitate was washed with 70% ethanol and resolved in 50 µl of TE buffer.

Total RNAs were extracted from the inner organs, intestines, muscles, and skin using the chaotropic extraction protocol for mouse pancreatic RNA, described by Robert [13]. Frozen tissues (3 - 5 mm^3^, 0.1 - 0.3 g) were submerged in 10 ml of TRIZOL Reagent (Life Technologies, Carlsbad, CA, USA), an amount three times larger than that recommended by the supplier. The tissues were then homogenized at 14,000 rpm three times for 30 s each using the Polytron homogenizer. After incubation for 5 min at room temperature, 2 ml of chloroform was added and the mixture was vortexed for 15 s. The mixture was incubated for 3 min at room temperature and centrifuged at 12,000 *g* for 10 min at 4°C. The upper aqueous phase (4 ml) was transferred to a fresh 50-ml tube and mixed with an equivalent amount of isopropyl alcohol. The sample was centrifuged at 12,000 *g* for 10 min and an RNA pellet was obtained. The RNA pellet was vortexed with 10 ml of 75% ethanol and centrifuged at 7500 *g* for 5 min at 4°C. The RNA pellet was air-dried and dissolved in 200 µl of RNase-free water by incubating for 10 min at 55°C. Further purification to remove contaminated genomic DNA was performed using a Monarch Total RNA Miniprep Kit (New England Biolabs, MA, USA), according to the supplier’s protocol. The RNA was eluted from a column by 100 µl of RNase-free water and kept at −80°C. The concentration and purity of the isolated RNA was determined using a spectrophotometer and denaturing agarose gel electrophoresis. The concentrations of RNA isolated from 11 tissue segments of inner organs, intestines, muscle, skin, and head ranged from 0.52-3.34 µg/µl or 0.38-1.60 µg/mg tissue.

### Determination of mitochondrial 16S/12S rRNA sequence

Primers (16SA-L-frog: 5’-CGCCTGTTTACCAAAAACAT-3’, 16SB-H-frog: 5’-CCGGTCTGAACTCAGATCACGT-3’, 12SB-F: 5’-AAACTGGGATTAGATACCCCACTAT-3’, 12SB-H: 5’-GAGGGTGACGGGCGGTGTGT-3’) used to amplify mitochondrial 16S or 12S rRNA partial regions were identical to those used in phylogenetic classification of Anura, reported by Hirota [14]. Polymerase chain reaction (PCR) was performed in 25 µl of 1x Q5 reaction buffer (0.5 units of Q5 High-Fidelity DNA Polymerase (New England Biolabs), 200 µM dNTPs, 0.5 µM forward primer, 0.5 µM reverse primer and 6.5 ng genomic DNA) using the thermal cycle program (30 s at 98°C, 30 cycles [30 s, 98°C; 30 s, 62°C; 30 s, 72°C], and 2 min at 72°C). PCR products were examined by agarose gel electrophoresis and sequenced using the same primers.

### mRNA library preparation and Illumina next-generation sequencing

An equivalent amount of total RNAs (0.52 - 3.34 µg/µl) isolated from 11 tissue segments of inner organs (3 segments), intestines (2 segments), muscles (2 segments), skin (3 segments), and head (1 segment) were mixed to obtain 100 ng/µl total RNA. The mixed total RNA was analyzed by Tape Station (RNA Screen tape, Agilent Technologies Ltd., USA) and determined to RIN (RNA integrity number) = 9.0. The poly (A)+ fraction was isolated from the total RNA, followed by its fragmentation. The fragmented mRNA was then reverse-transcribed to double-stranded cDNA using random hexamers. The double-stranded cDNA fragments were processed for adaptor ligation, size selection for 200-bp inserts, and amplification of strand-specific cDNA libraries. Prepared libraries were subjected to paired-end 2 x 100 bp sequencing on the HiSeq 2500 platform with v4 chemistry.

### *De novo* transcriptome assembly and bioinformatic analysis

All analyses were performed mainly using the CLC Genomics Workbench 11.0 (QIAGEN Bioinformatics, Germany). The 2 x 100 bp paired-end reads were checked in terms of the sequencing quality and trimmed (removal of adaptor and duplication) with quality score limit of 0.05 and a maximum number of two ambiguous nucleotides. The clean reads were then assembled and aligned with a word size of 20, bubble size of 50, and minimum transcript length of 200, based on the algorithm of the De Brujin graph. The open reading frame (ORF) in the assembled *P. adspersus* transcripts was predicted under the following conditions: start codon as ATG, search as Both Strand, open-ended sequence, minimum length (codons) as 25, and genetic code as standard. The transcripts were homology searched using local BLASTx (National Center for Biotechnology Information, NCBI) against the Uniprot database (https://www.uniprot.org/) (all-proteins or *Xenopus tropicalis*) or SmProt database (http://bioinfo.ibp.ac.cn/SmProt/index.htm) (small proteins curated from literature mining, known databases, ribosome profiling data, or MS data). Homologous proteins found in the Uniprot database (*X. tropicalis*) with an E-value lower than 1E-5 were subjected to gene ontology (GO) analysis (https://www.uniprot.org) [15] to assign the GO terms of biologic processes, molecular functions, and cellular components.

### Data availability

The data sets supporting the results of this article are available from the following databases.

Mitochondrial 16S rRNA of the African bullfrog (583 bp): GenBank: LC440402 Mitochondrial 12S rRNA of the African bullfrog (424 bp): GenBank: LC440403 Cytochrome *b* of the African bullfrog (transcripts #275): GenBank: LC440404 Raw reads of 2 x 100 bp RNA sequence of the African bullfrog: DDBJ sequencing read archive (SRA): DRR164258, BioProject: PRJDB7457, BioSample: SAMD00143251, Experiment: DRX154877 Transcript sequences of the African bullfrog: GenBank: ICLD01000001-ICLD01101682

## Results

### Confirmation of animal species used in this study

To confirm that the experimental animal used in this study was in fact the African bullfrog, a captive bred frog was raised for 5 years and the appearance of the adult frog was observed. The frog was approximately 20 cm in length and had olive-green skin with longitudinal skin folds (**Fig 1A**). These morphologic characteristics were consistent with those of the African bullfrog reported by Channing [16] and Bishop [17], and clearly differed from the characteristics of *P. edulis*, a species closely related to the African bullfrog that is 12 cm in length and has olive green-yellow skin with spotted folds. Furthermore, we determined the nucleotide sequence of mitochondrial 16S and 12S rRNA of the frog used for subsequent *de novo* assembly analysis. NCBI BLAST analysis revealed that the mitochondrial 16S rRNA (583 bp, GenBank: LC440402) was 100% identical to the mitochondrial 16S rRNA of *Pyxicephalus cf. adspersus* FB-2006 isolate 0954 (551 bp, GenBank: DQ347304) (**Fig 1B**). The mitochondrial 12S rRNA (424 bp, GenBank: LC440403) was 99% identical to the mitochondrial 12S rRNA of *P. cf. adspersus* FB-2006 isolate 0954 (1,364 bp, GenBank: DQ347013) (**S1 Fig**). These morphologic characteristics and the results of the homology analysis of rRNA sequences confirmed that the frog used in this study was *P. adspersus*, the African bullfrog.

**Fig 1.**
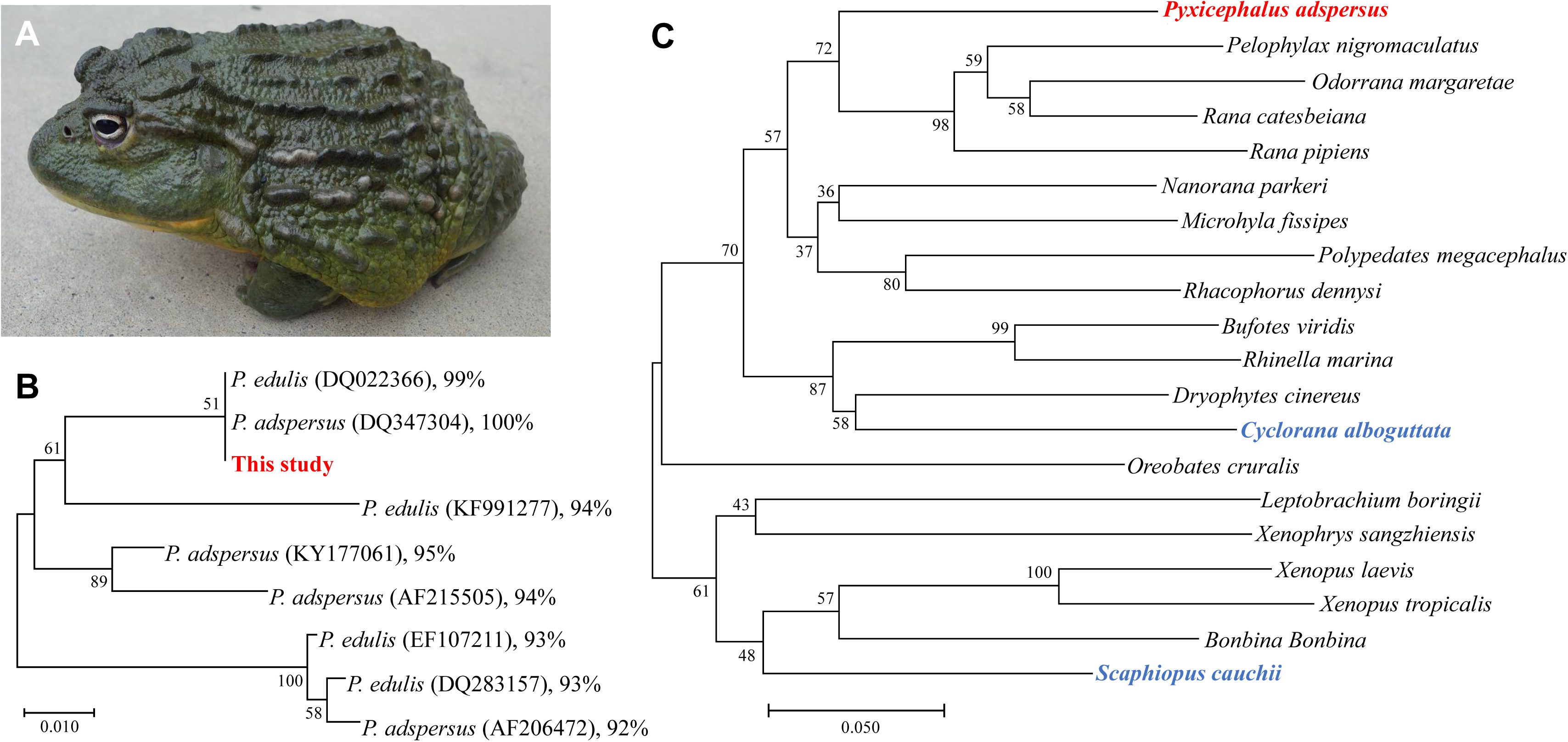
The African bullfrog used in this study. A) The adult male African bullfrog used in this study is shown. B) Phylogenetic tree analysis of the 16S rRNA sequence between the frog used in this study and several *P. adspersus* and *P. edulis* species deposited in the database. Sequences were aligned using MUSCLE algorithm in MEGA X. Phylogenetic tree was constructed using neighbor-joining algorithm and the node support values were assessed by bootstrap test of 1000 replicates. C)The phylogenetic tree using the 16S rRNA sequence between the representative frog species belonging to the major family of Anura. *P. adspersus* used in this study is shown in red and that of the other two aestivating frogs are shown in blue. Phylogenetic tree was constructed in the same manner in Fig 1B.

The two types of frog, *P. adspersus* and *P. edulis*, are sometimes confused with each other [16,18]. Top hits against the 16S and 12S rRNA sequences of the frog used in this study are found in both species in no particular order. Furthermore, the phylogenetic tree analysis of the top hits 16S and 12S rRNA sequences revealed that the two species were not clustered (**Fig 1B and S1 Fig**). We speculate that the nomenclatures of these rRNA sequences were mistakenly registered in the database.

Figure 1C shows the phylogenetic tree of mitochondrial 16S rRNA partial sequences of the African bullfrog used in this study and of representative frog species belonging to the major family of Anura. The two aestivating frogs, *C. alboguttata* and *S. cauchii*, are phylogenetically distant from *P. adspersus*, as suggested by Pyron’s report [19].

### RNA sequencing and *de novo* assembly

RNA sequencing and *de novo* assembly data are summarized in Table 1. RNA sequencing produced 75,604,146 raw reads. *De novo* transcriptome assembly of the trimmed 75,320,390 reads generated 101,682 transcripts. The transcript size ranged from 236 to 35,997 bases with a mean length of 866 bases and an N50 length of 1,701 bases. The number of transcripts ranging from 1000 to 1999 bases was highest among that of transcripts ranging from 500 to 5,000 bases (**Fig 2**).

**Table 1.** Summary of the RNA sequence reads and the *de novo* transcriptome assembly for the African bullfrog.

|  |  |  |
| --- | --- | --- |
| **Sequencing results** | Length of raw reads (bp) | 100 |
|  | Total number of raw reads | 75,604,146 |
|  | Total number of clean reads | 75,320,390 |
| ***De novo* transcriptome assembly results** | Total number of transcripts | 101,682 |
|  | Total length of all transcripts (bases) | 88,093,202 |
|  | GC contents of all transcripts (%) | 40.8 |
|  | N50 length of all transcripts (bases) | 1,701 |
|  | Mean length | 866 |

**Fig 2.**
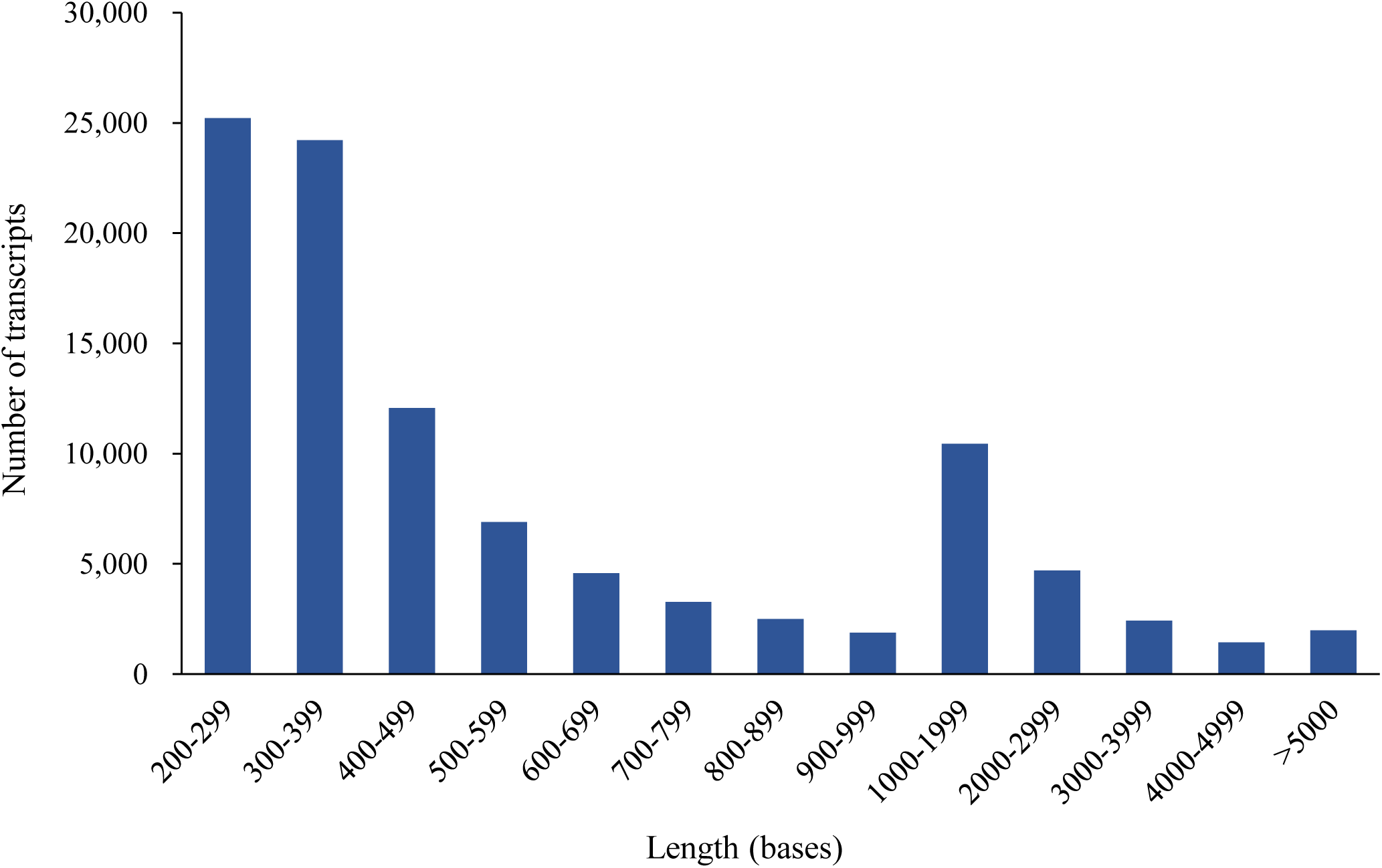
Distribution of the length of the African bullfrog transcripts. The number of transcripts was count and the transcript length was determined.

*In silico* analysis to identify ORF-encoding proteins larger than 25 amino acids revealed that 100,093 transcripts (98.4% of total transcripts) were protein-coding genes (**Table 2**). The longest ORF-encoded protein was 11,679 amino acids long. The average ORF-encoded protein was 140.7 amino acids long. In contrast, 1590 transcripts (1.6% of total transcripts) had no ORF, suggesting that these transcripts were non-coding RNAs. When the ORF detection conditions were changed to 30, 50 or 100 amino acids long, the number of transcripts carrying the ORF was decreased to 97,122 (95.5%), 71,305 (70.1%) or 29,296 (28.8%), respectively (**Table 2**).

**Table 2.** Detection of ORF in the African bullfrog transcripts.

|  | Minimum ORF length |  |  |  |
| --- | --- | --- | --- | --- |
|  | 25 aa | 30 aa | 50 aa | 100 aa |
| Total transcripts | 101,683 |  |  |  |
| Detected ORF | 729,832 | 562,570 | 219,579 | 45,086 |
| Transcripts coding ORF | 100,093<br>(98.4%) | 97,122<br>(95.5%) | 71,305<br>(70.1%) | 29,296<br>(28.8%) |
| Maximum ORF length | 11,679 | 11,679 | 11,679 | 11,679 |
| Minimum ORF length | 25 | 30 | 50 | 100 |
| Average ORF length (aa) | 140.7 | 144.1 | 181.4 | 341.2 |
| Transcripts of non-coding RNA | 1,590<br>(1.6%) | 4,561<br>(4.6%) | 30,378<br>(29.9%) | 72,387<br>(71.2%) |

Within these *de novo* assembled transcripts, we identified that transcript #275 had high similarity with cytochrome *b* on the basis of the BLASTx search. The DNA sequence of transcript #275 (GenBank: LC440404) was 100% identical to *P. adspersus* isolate Brak01 cytochrome *b* gene (GenBank: FJ613662) (**S2 Fig**). This result confirmed that that the frog used in this study was *P. adspersus*.

### Transcriptome annotation

We performed a BLASTx homology search of the African bullfrog transcripts against the Uniprot protein database (all-proteins or *Xenopus tropicalis* proteins). In the all-proteins database, 27,286 transcripts (26.8%) had an E-value lower than 1E-5, 53,064 transcripts (52.2%) had an E-value higher than 1E-5, and 21,332 transcripts (21.0%) showed no hits with the database (**Fig 3A**, All proteins). In the *X. tropicalis* protein database, 28,572 transcripts (28.1%) had an E-value lower than 1E-5, 64,963 transcripts (63.9%) had an E-value higher than 1E-5, and 8,147 transcripts (8.0%) showed no hits with the database (**Fig 3A**, *X. tropicalis*). Figure 3B shows the distribution of transcripts with an E-value lower than 1E-5 according to the amino acid identity with the protein database. 22,670 transcripts (83.1%) and 25,407 transcripts (88.9%) had an amino acid identity higher than 50% in the all-proteins database or in the *X. tropicalis* protein database, respectively.

**Fig 3.**
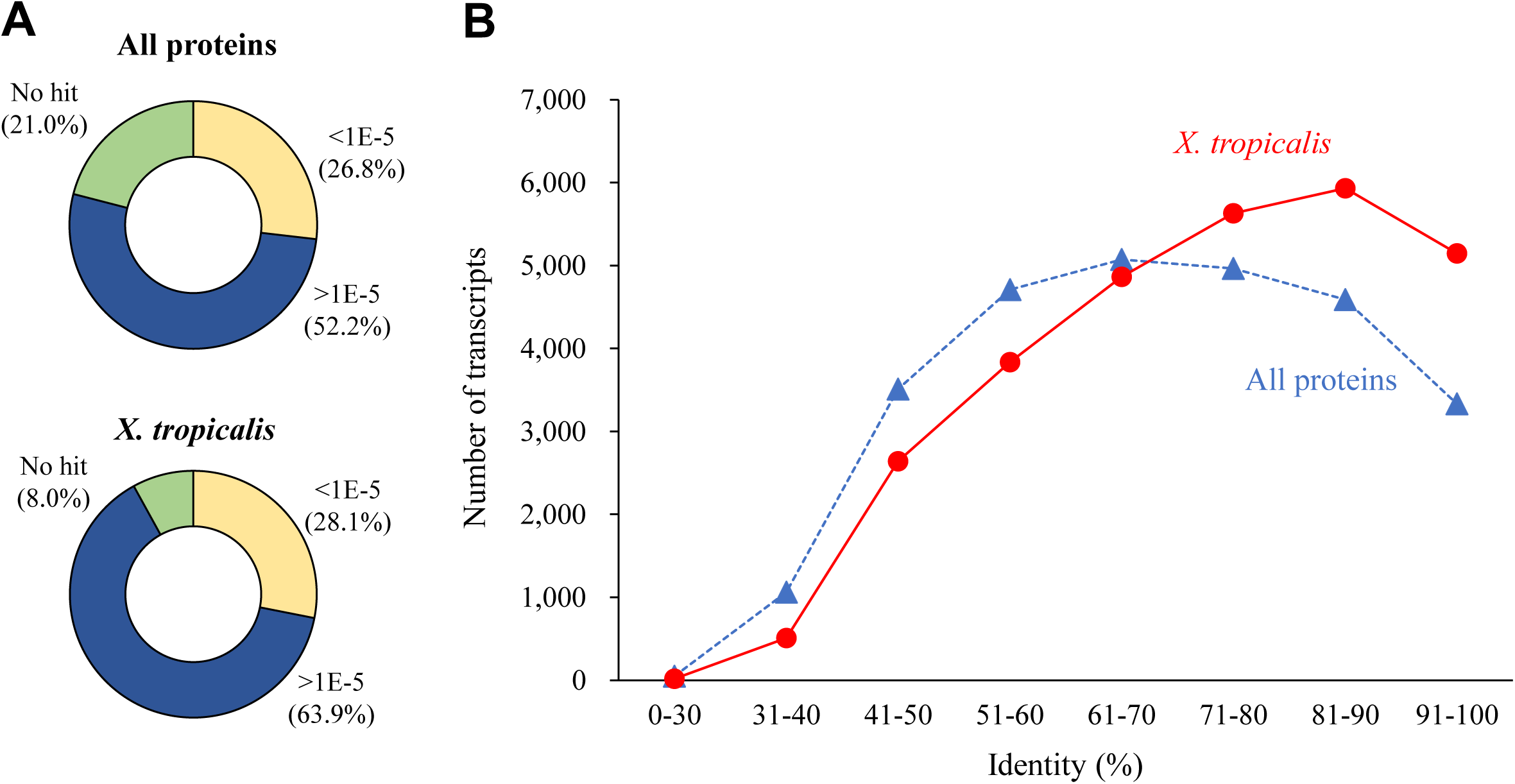
Similarity of the African bullfrog transcripts against the Uniprot protein databases. A) BLASTx search of the African bullfrog transcripts against the Uniprot database (all-proteins or *X. tropicalis* proteins), and the number of transcripts were counted according to E-values. B) Distribution of the identity of the African bullfrog transcripts with an E-value lower than 1E-05 against the *X. tropicalis* protein database.

To identify small proteins that are absent in the Uniprot database, we performed a BLASTx homology search of the African bullfrog transcripts against the SmProt small protein databases (small proteins curated from literature mining, known databases, ribosome profiling data, or MS data). In any of the SmProt small protein databases, more than 80% transcripts had an E-value higher than 1E-5 or showed no hit (**Fig 4A**). We further examined whether the African bullfrog transcripts having little similarity to the Uniprot all protein database (53,064 transcripts, >1E-5; 21332 transcripts, no hits) and the Uniprot X. tropicalis protein database (64,963 transcripts, >1E-5; 8,147 transcripts, no hits) shows little similarity against the SmProt databases. As a result, more than 99% non-annotated transcripts in the Uniprot all protein database as well as those in the Uniprot X. tropicalis protein database had E-value higher than 1E-5 or no hits in the SmProt databases (**Fig 4B**).

**Fig 4.**
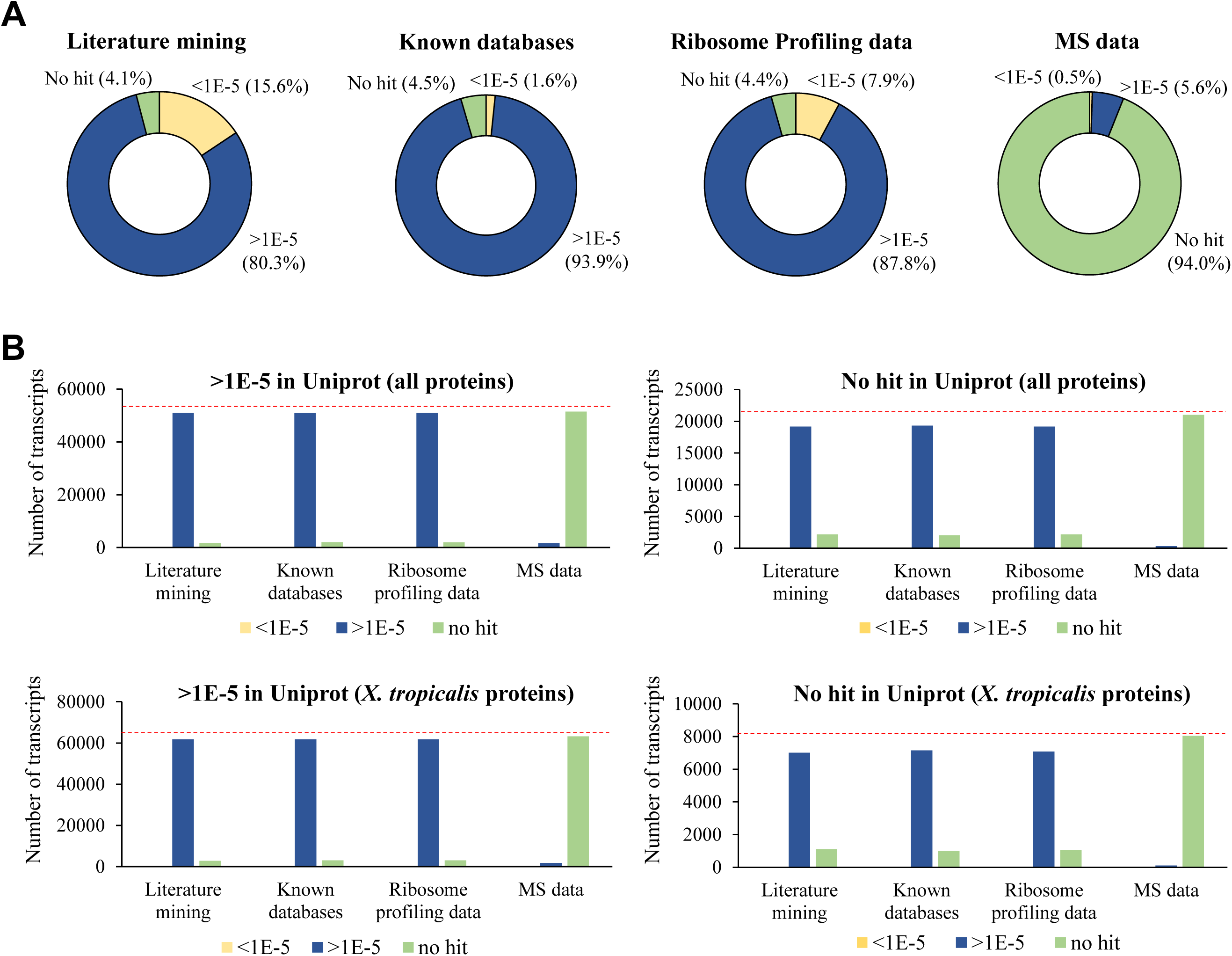
Similarity of the African bullfrog transcripts against the SmProt small protein databases. A) BLASTx search of the African bullfrog transcripts against the SmProt databases including literature mining, ribosome profiling data, known databases, and MS data. The number of transcripts were counted according to E-values. B) The African bullfrog 74,396 transcripts having little similarity to Uniprot all protein database (53,064 transcripts, >1E-5; 21,332 transcripts, no hit) were subjected to BLASTx search against SmProt database (Literature mining, Known database, Ribosome profiling, and MS data). Red dotted line represents the total number of transcripts subjected to this analysis.

### Gene ontology analysis

The 28,572 transcripts with an E-value lower than 1E-5 in Uniprot *X. tropicalis* protein database were subjected to gene ontology (GO) analysis and classified according to three major GO categories: cellular components, molecular function, and biologic processes (**Fig 5**). In the cellular component category, 11,634 (33.4%) and 7,689 (22.0%) genes were associated with the cellular components and membrane components, respectively. In the molecular function category, 11,142 (45.6%) and 6,995 (28.6%) genes were associated with binding and catalytic activity, respectively. In the biologic process category, 9,765 (30.5%), 6,213 (19.4%), and 5,553 (17.3%) genes were associated with cellular processes, biologic regulation, and metabolic processes, respectively.

**Fig 5.**
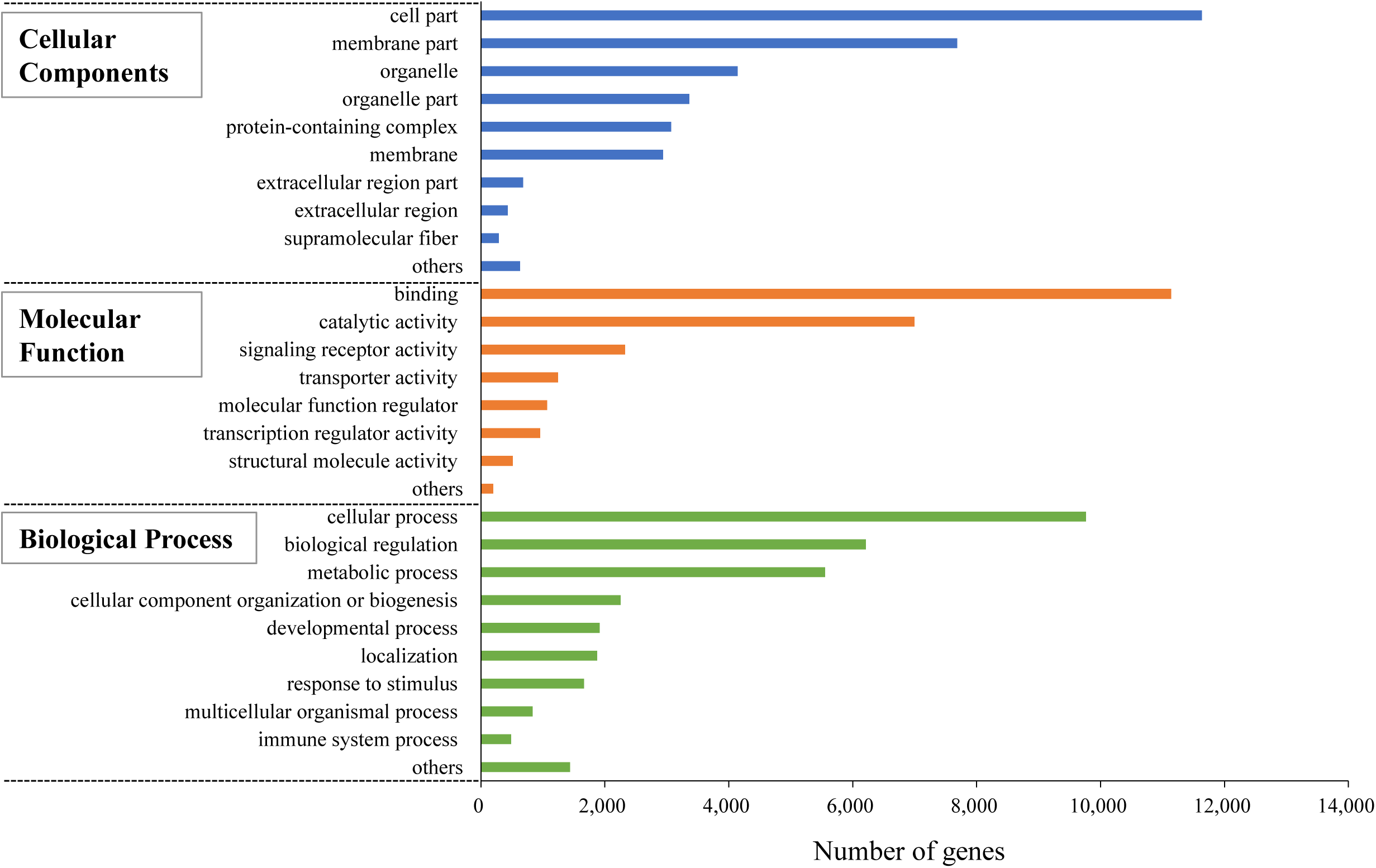
Gene ontology terms assigned to the African bullfrog genes. The 28,570 genes with high identity (E-value lower than 1E-5) in a BLASTx search of the *X. tropicalis* protein database were subjected to gene ontology analysis.

### Comparison with the aestivation-related genes reported in other organisms

To determine whether the African bullfrog transcripts contain genes homologous to the aestivation-related genes of the two aestivating frogs and the African lungfish, we performed a homology search of aestivation-related genes against the African bullfrog transcripts (**Table 3**) [8,9,10]. The #2083 transcript in the African bullfrog had high identity with riboflavin-binding protein precursor (RfBP) in Couch’s spadefoot toad, which has increased expression during aestivation and might be involved in egg maturation in the frog during the aestivating stage. The African bullfrog genes had high identity with genes of the Green striped burrowing frog, which are highly upregulated during aestivation, such as adaptor related protein complex 3 (beta 2 subunit) that functions in vesicle budding from the Golgi apparatus; an immunoglobulin heavy chain that recognizes foreign antigens; and a serine/threonine protein kinase, Chk1, that activates DNA repair in response to DNA damage. In addition, the African bullfrog genes had high identity with African lungfish genes that are expressed during aestivation, such as argininosuccinate synthetase 1 and carbamoylphosphate synthetase III, which are key enzymes in the ornithine-urea cycle and function to detoxify ammonia to urea; rhesus family C glycoprotein that maintains muscle function; and superoxide dismutase 1, an anti-oxidant. These findings suggest that African bullfrog has a set of genes that are highly expressed in other vertebrates during aestivation.

**Table 3.** Presence of aestivation-related genes in the African Bullfrog.

| Species | Gene | Predicted Function | Top Hit Contig of African Bullfrog | E-value | % Identity |
| --- | --- | --- | --- | --- | --- |
| *Scaphiopus couchii*<br>(Couch's spadefoot toad) | Riboflavin binding protein precursor<br>(AF102545) | Maturation of eggs in preparation for the explosive breeding | #2083<br>(ICLD012083) | 8.63E-114 | 67 |
| *Cyclorana alboguttata*<br>(Green-striped burrowing frog) | Adaptor related protein complex 3, beta 2 subunit<br>(contig #8496) | Budding of vesicles from the golgi membrane | #4968<br>(ICLD014968) | 0 | 100 |
|  | Solute carrier family 28 member 3-like<br>(contig #7653) | Homeostasis of endogenous nucleosides | #2599<br>(ICLD012599) | 1.99E-45 | 100 |
|  | Nuclease HARBI1-like<br>(contig #14756) | Nuclease activity | #59155<br>(ICLD0159155) | 1.31E-122 | 88 |
|  | Ribosome biogenesis protein NSA2-like<br>(contig #40) | Biogenesis of the 60S ribosomal subunit | #4109<br>(ICLD014109) | 5.97E-103 | 99 |
|  | Coiled coil domain containing protein 81<br>(contig #15484) | Positive regulator of T-cell maturation and inflammatory function | #18527<br>(ICLD0118527) | 0 | 75 |
|  | Pentraxin precursor<br>(contig #27476) | Mediating uptake of synaptic material | #3473<br>(ICLD013473) | 7.25E-171 | 90 |
|  | Immunoglobulin heavy prechain<br>(contig #36672) | Recognition of foreign antigens and initiation of immune responses | #3055<br>(ICLD013055) | 1.01E-118 | 82 |
|  | Receptor type tyrosine protein phosphatase T-like<br>(contig #7294) | Signal transduction and cellular adhesion | #10027<br>(ICLD0110027) | 0 | 99 |
|  | Immunoglobulin J chain precursor<br>(contig #48432) | Serving to link two monomer units of either IgM or IgA | #1190<br>(ICLD011190) | 1.82E-15 | 76 |
|  | UHRF1 binding protein 1-like<br>(contig #1661) | Negative regulator of cell growth | #8912<br>(ICLD018912) | 0 | 95 |
|  | Inositol 3 phosphate synthase 1<br>(contig #6523) | Key enzyme in myo-inositol biosynthesis pathway that catalyzes the conversion of glucose 6-phosphate to 1-myo-inositol 1-phosphate | #13219<br>(ICLD0113219) | 0 | 93 |
|  | Serine/threonine protein kinase Chk1<br>(contig #13899) | Activation of DNA repair in response to<br>the presence of DNA damage or<br>unreplicated DNA | #24942<br>(ICLD0124942) | 0 | 85 |
| *Protopterus annectens*<br>(African Lungfish) | Rhesus family C glycoprotein<br>(KX583647) | Glycolysis and muscle development | #10697<br>(ICLD0110697) | 4.05E-162 | 79 |
|  | Argininosuccinate synthetase 1<br>(JZ575533) | Ornithine-urea cycle capacity to detoxify<br>ammonia to urea | #297<br>(ICLD01297) | 1.72E-56 | 81 |
|  | Carbamoylphosphate synthetase III<br>(JZ575539) | Ornithine-urea cycle capacity to detoxify<br>ammonia to urea | #265<br>(ICLD01265) | 3.70E-108 | 91 |
|  | Betain homocysteine-S-transferase 1<br>(JZ575536) | Prevent the accumulation of hepatic<br>homocysteine | #243<br>(ICLD01243) | 5.19E-86 | 94 |
|  | Superoxide dismutase 1<br>(JZ575606) | Anti ROX | #1087<br>(ICLD011087) | 2.73E-62 | 100 |

tBLASTx search was performed to identify the African bullfrog contig having high identity to the genes that are up-regulated during the aestivation
in the three organisms. The ID numbers in parenthesis indicate Genbank accession numbers or contig numbers (*C. alboguttata*).

## Discussion

The African bullfrog is well known as an aestivating frog and is bred all over the world. Molecular biologic analysis of this frog, however, has not been performed and no genetic information has been available. In this study, we performed the first *de novo* transcriptome assembly and identified 101,682 transcripts. Of the 101,682 transcripts identified, 53,064 (52.2%) had an E-value higher than 1E-5 and 21,332 transcripts (21.0%) had no hits in a BLASTx search in the Uniprot all protein database. In addition, 64,963 (63.9%) had an E-value higher than 1E-5 and 8,147 transcripts (8.0%) had no hits in the Uniprot *X. tropicalis* protein database. These results suggest that more than 70% of total transcripts are novel genes in the African bullfrog. Of these novel genes, more than 99% had little similarity to the SmProt small protein database. The result suggests that the high proportion of novel genes of the African bullfrog are not explained by the genes coding small proteins, which are available in the database. The high proportion of the novel genes in total transcripts is also observed in other organisms whose *de novo* transcriptome assembly was recently performed, such as the Caecilian (*Caecilia tentaculate*), the Sodom apple (*Calotropis gigantea*), and the Black ant (*Formica fusca*) [20, 21, 22], and confirms that *de novo* transcriptome assembly is a powerful tool to identify novel genes. The novel genes of the African bullfrog might encode proteins without known domains, or non-coding RNAs. With regard to the very limited genomic and transcriptomic information in amphibians, the novel genes of the African bullfrog are the significant biologic resources to perform molecular biologic analysis of the physiologic functions of the African bullfrog, including its aestivation ability.

In contrast, 28,572 transcripts (29.1%) had an E-value lower than 1E-5 in the *X. tropicalis* database, suggesting that these genes are conserved in *X. tropicalis*. These genes were subjected to gene ontology analysis and their molecular functions were estimated. They are assumed to have functions conserved between *P. adspersus* and *X. tropicalis*. Furthermore, we revealed that the African bullfrog has a set of aestivation-related genes, which has been reported in two aestivating frogs and the African lungfish. These genes presumably function in the aestivation mechanism conserved between different aestivating vertebrates. The aestivation-related genes are not specific to the aestivating organisms, but are conserved in other non-aestivating organisms, and must therefore be transcriptionally controlled by factor(s) specific to the aestivating environment or the aestivating organisms. How the conserved genes are regulated during aestivation remains an important unanswered question.

The genetic information of the African bullfrog obtained in this study enables future comparative transcriptome analysis between the active stage and the aestivating stage of the frog and contributes to uncover the molecular mechanisms of aestivation.

## Supporting Information

**S1 Fig. The phylogenetic tree analysis of 12S rRNA sequence between the frog used in this study and several *P. adspersus* and *P. edulis* species deposited in database**. Sequences were aligned using MUSCLE algorithm implemented in MEGA X. Phylogenetic tree was constructed using neighbor-joining algorithm with bootstrap test of 1000 replicates.

**S2 Fig. The phylogenetic tree analysis of cytochrome b partial sequence between the frog used in this study and several frogs deposited in database**. Sequence of cytochrome *b* of the African bullfrog (GenBank: LC440404) was derived from *de novo* assembled transcript #275. Sequences were aligned using MUSCLE algorithm implemented in MEGA X. Phylogenetic tree was constructed using neighbor-joining algorithm with bootstrap test of 1000 replicates.

