## Supplementary figures and images for "*De novo* transcriptome assembly of the African bullfrog *Pyxicephalus adspersus* for molecular analysis of aestivation"

### Supplemental Figure 1

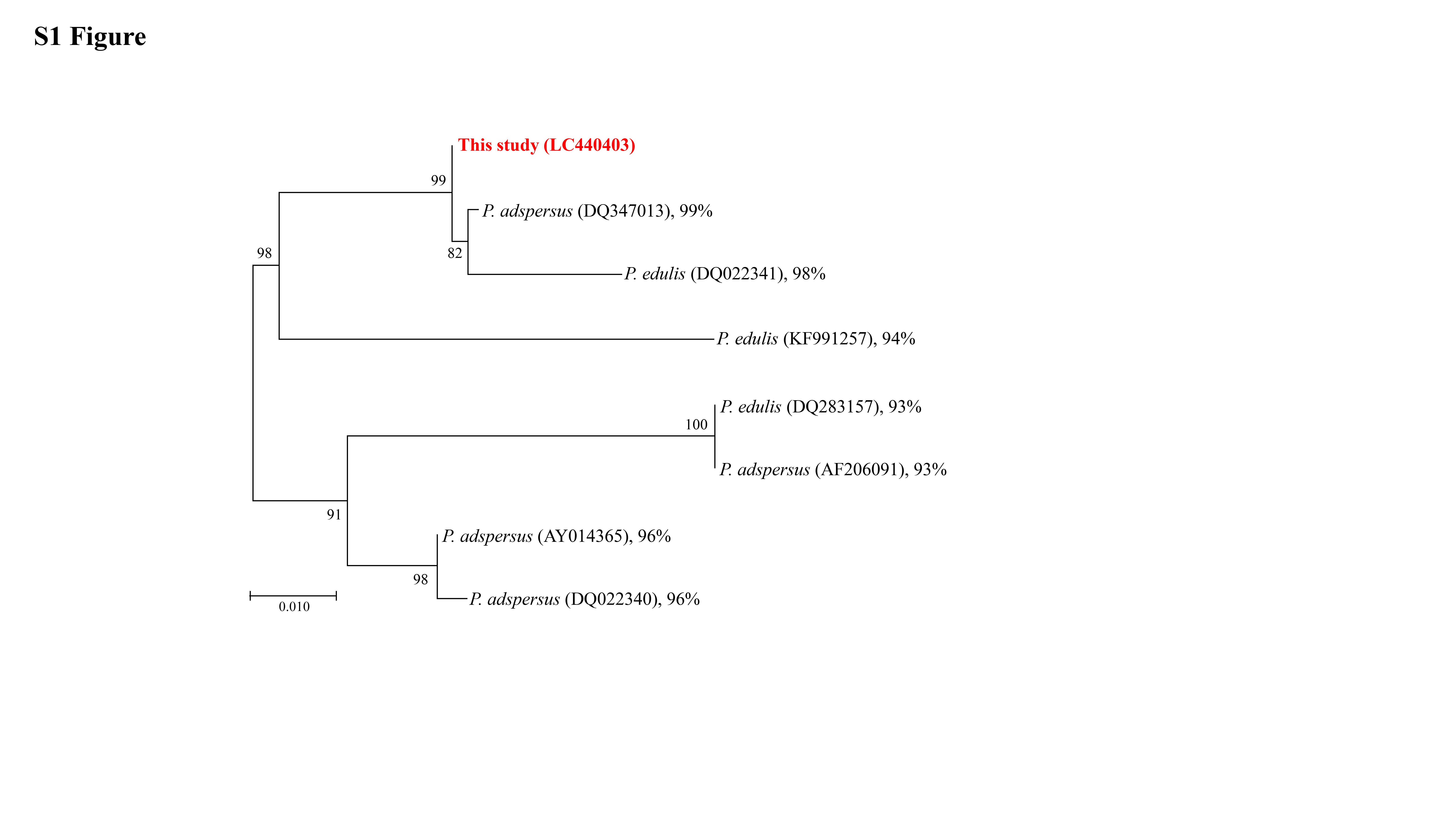

### Supplemental Figure 2

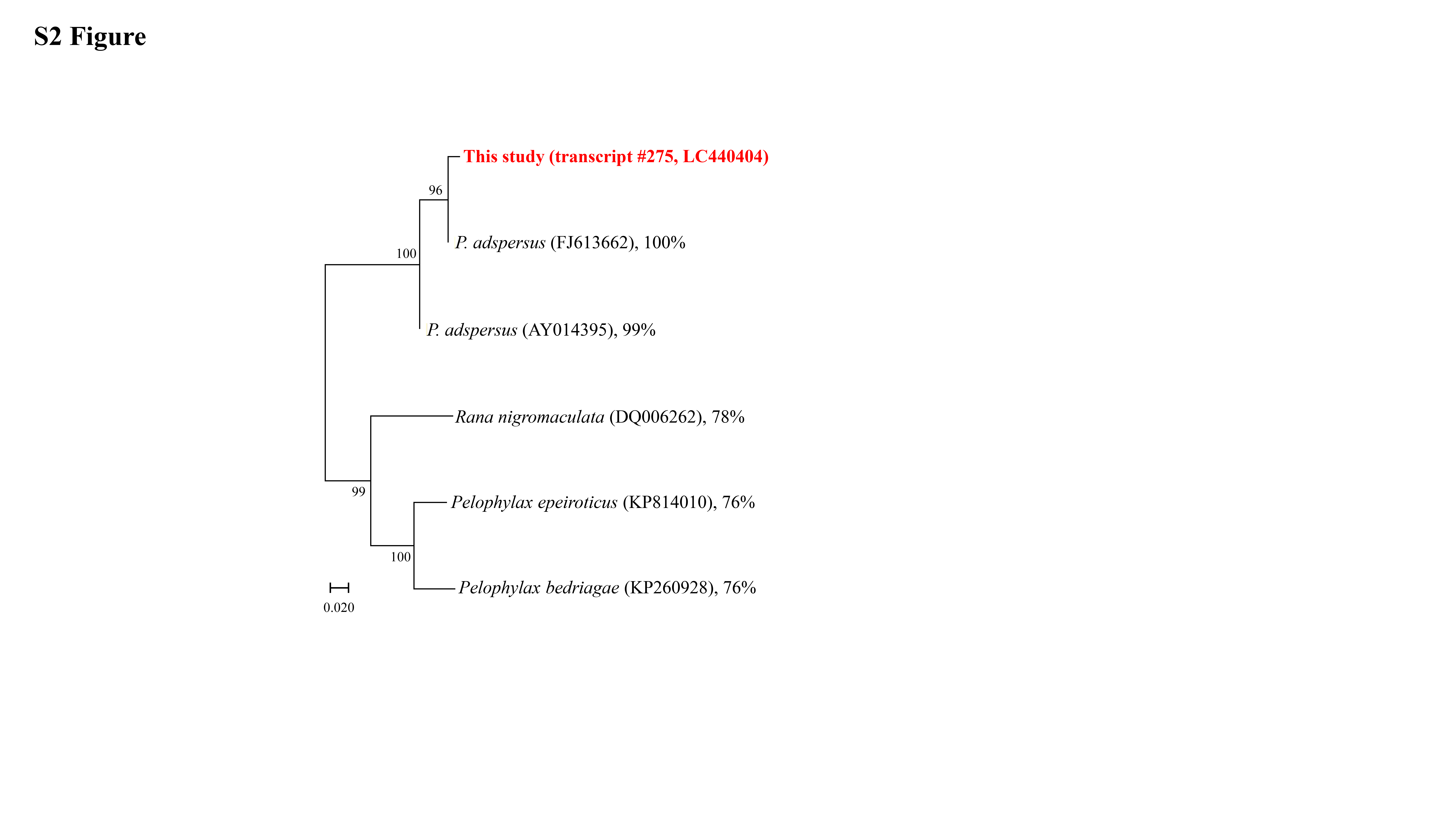
